# Natural bacterial assemblages in *Arabidopsis thaliana* tissues become more distinguishable and diverse during host development

**DOI:** 10.1101/2020.03.04.958165

**Authors:** Kathleen Beilsmith, Matthew Perisin, Joy Bergelson

## Abstract

To study the spatial and temporal dynamics of bacterial colonization under field conditions, we planted and sampled *Arabidopsis thaliana* during two years at two Michigan sites and surveyed colonists by sequencing 16S rRNA gene amplicons. Mosaic and dynamic assemblages revealed the plant as a patchwork of tissue habitats that differentiated with age. Although assemblages primarily varied between roots and shoots, amplicon sequence variants (ASVs) also differentiated phyllosphere tissues. Increasing assemblage diversity indicated that variants dispersed more widely over time, decreasing the importance of stochastic variation in early colonization relative to tissue differences. As tissues underwent developmental transitions, the root and phyllosphere assemblages became more distinct. This pattern was driven by common variants rather than those restricted to a particular tissue or transiently present at one developmental stage. Patterns also depended critically on fine phylogenetic resolution: when ASVs were grouped at coarse taxonomic levels, their associations with host tissue and age weakened. Thus, the observed spatial and temporal variation in colonization depended upon bacterial traits that were not broadly shared at the family level. Some colonists were consistently more successful at entering specific tissues, as evidenced by their repeatable spatial prevalence distributions across sites and years. However, these variants did not overtake plant assemblages, which instead became more even over time. Together, these results suggested that the increasing effect of tissue type was related to colonization bottlenecks for specific ASVs rather than to their ability to dominate other colonists once established.

**Importance:** Developing synthetic microbial communities that can increase plant yield or deter pathogens requires basic research on several fronts, including the efficiency with which microbes colonize plant tissues, how plant genes shape the microbiome, and the microbe-microbe interactions involved in community assembly. Findings on each of these fronts depend upon the spatial and temporal scales at which plant microbiomes are surveyed. In our study, phyllosphere tissues housed increasingly distinct microbial assemblages as plants aged, indicating that plants can be considered as collections of tissue habitats in which microbial colonists-- natural or synthetic-- establish with differing success. Relationships between host genes and community diversity might vary depending on when samples are collected, given that assemblages grew more diverse as plants aged. Both spatial and temporal trends weakened when colonists were grouped by family, suggesting that functional rather than taxonomic profiling will be necessary to understand the basis for differences in colonization success.

## INTRODUCTION

As plant tissues emerge and grow, new habitats are created for microbial colonists (1, 2). While they represent only a fraction of colonist diversity (3, 4), bacterial endophytes are important to host plant fitness because of their potential to affect nutrient uptake (5, 6, 7), stress responses (8), and defenses against pathogens (9, 10). Given these activities, natural and engineered bacterial communities have been proposed as tools for sustainably enhancing plant growth and stress resistance (11). When lineages in these communities are pathogenic or beneficial to the host, the efficiency with which they enter plant tissue is of particular interest (12, 13, 14). Despite stochasticity in colonization (15, 16, 17), there is evidence that plants selectively filter the bacteria colonizing the intercellular space in their tissues. Characterizing spatial and temporal variation in this filtering is key to understanding how natural and cultivated communities assemble in the endosphere.

The idea that plant tissue filters bacterial colonists is supported by the observation that endophytic communities display only a fraction of the diversity found in soil. For example, the diversity of taxa found in the root endosphere is lower than in rhizosphere soil (18). Furthermore, a subset of bacterial families is found at higher relative abundance in roots than in soil (19). Filtering is likely due in part to differences between soil and root cell walls as substrates for colonization, as indicated by similarities between communities in live roots and wood slivers exposed to the same field-collected soil inocula (20). Filtering by living tissue may also involve selection for or against specific bacterial lineages, as indicated by community members enriched over soil levels in roots but not in wood sliver samples and vice versa (20).

Although plant tissue appears to filter bacterial colonists, we do not yet fully understand how spatial and temporal variation influences this process in natural environments. Variation in abiotic factors and the pool of soil colonists at planting sites can influence the efficiency with which bacterial lineages enter plants, leading to associations between geographic location and the composition of bacterial assemblages in plant tissues (21, 22, 23). Within plants, recent evidence suggests that communities in different tissues are composed of a common pool of systemic colonists (24, 25). However, individual lineages can display differences in colonization efficiency between roots and stems (26) and several studies report variation between the bacterial assemblages found in different tissues (27, 28, 29), supporting the idea that some bacteria are more successful than others in colonizing a given habitat within the plant. Variation in assemblage composition is also observed when plant tissues are sampled at different developmental stages (22, 30, 31). These temporal trends could be related to the time available for bacterial colonization of plant tissues before sampling, changes in how hosts filter colonists with age, or interactions arising as more bacteria cooperate or compete within the host. Since bacteria alter plant tissues upon arrival, the host response to established colonizers could also change the efficacy of colonization later in development (32, 33).

Since most surveys of plant colonists have either focused on a single tissue or taken only a snapshot of community composition in time, it is difficult to compare the extent and interaction of geographic, tissue-level, and temporal effects on plant endophyte filtering. Adding to the body of work exploring host plant control of colonization, we compared the relative influence of plant tissue type, age, harvest site, and year on the bacteria that naturally colonized a common haplotype of an annual plant, *Arabidopsis thaliana*. To understand the distribution of the bacterial traits driving these patterns, we examined whether relationships between variables and assemblage composition depended on the taxonomic level at which the surveyed colonists were grouped. To ascertain the consistency of spatial and temporal colonization patterns, we compared the tissues and ages at which bacterial lineages reached maximum prevalence or abundance between sites and years. In addition, we characterized how the diversity and evenness of colonists changed across plant tissues throughout development.

## RESULTS

We planted surface-sterilized seeds of *A. thaliana* accessions from a single North American haplotype (Supplementary Table S1) at two southwest Michigan sites in two consecutive years. These accessions germinate in the fall, overwinter as small rosettes, flower in the spring, and senesce in the early summer. We harvested roots and rosette leaves throughout vegetative growth and also stems, cauline leaves, flowers, and siliques as they became available during flowering and senescence (Supplementary Table S2). To enrich for endophytes, bacteria were washed from the surface of plant tissues by repeated vortexing in surfactant buffer (Supplementary Text S1). Topsoil was also collected from field sites at each timepoint during the second year of study. Bacterial lineages in the soil and plant tissue samples were quantified by amplification and sequencing of the V5, V6, and V7 regions of the 16S rRNA gene (16S) (29). The 16S sequences were grouped into amplicon sequence variants (ASVs) with DADA2 (34) in QIIME2 (35). After filtering out singleton 16S variants and those from plant organelles, 10,803 ASVs were tallied for 1,272 samples (Table 1). A phylogenetic tree for the variants was inferred with FastTree using MAFFT-aligned 16S sequences (36, 37). The ASVs were classified at seven taxonomic ranks based on the SILVA 16S database (38).

**TABLE 1.** Samples in the study after quality control (at site ME, at site WW)

|  | Soil | Roots | Rosettes | Stems | C. Leaves | Flowers | Siliques | Total |
| --- | --- | --- | --- | --- | --- | --- | --- | --- |
| No Plant | 15, 19 |  |  |  |  |  |  | 34 |
| Two-Leaf Plant | 6, 4 | 3, 4 | 8, 10 |  |  |  |  | 35 |
| Four-Leaf Plant |  | 29, 8 | 27, 29 |  |  |  |  | 93 |
| Six-Leaf Plant | 8, 4 | 32, 18 | 45, 26 |  |  |  |  | 133 |
| Eight-Leaf Plant |  | 37, 24 | 41, 35 |  |  |  |  | 137 |
| Flowering Plant | 4, 3 | 61, 38 | 64, 49 | 41, 38 | 24, 22 | 42, 28 | 14, 25 | 453 |
| Senescent Plant | 6, 8 | 60, 64 |  | 54, 75 |  |  | 49, 71 | 387 |
| Total | 77 | 378 | 334 | 208 | 46 | 70 | 159 | 1272 |

### Bacterial assemblage composition was associated with plant tissue type and developmental stage

Plants shaped the bacterial assemblages they hosted, making them distinct from those in the surrounding soil (Supplementary Text S2). Rather than a single host environment, the plant appeared to be a collection of microbe habitats defined by tissue type and age. Samples from the same tissue or stage clearly shared a higher proportion of members than randomly compared samples (Figure 1). When samples from multiple tissues of the same individual plant were available, comparisons showed that they did not share a significantly higher proportion of members than randomly paired samples. Bacterial assemblages therefore appeared to be more similar between samples of the same tissue type from different plants than between samples from different tissue habitats in the same plant. Common environments also influenced assemblages, as evidenced by increased membership overlap in samples from the same site or year compared to random samples. Despite these patterns, the low proportion of members shared within groups conditioned on any study variable (< 15%) underscored the high variability of colonization.

**Figure 1.**
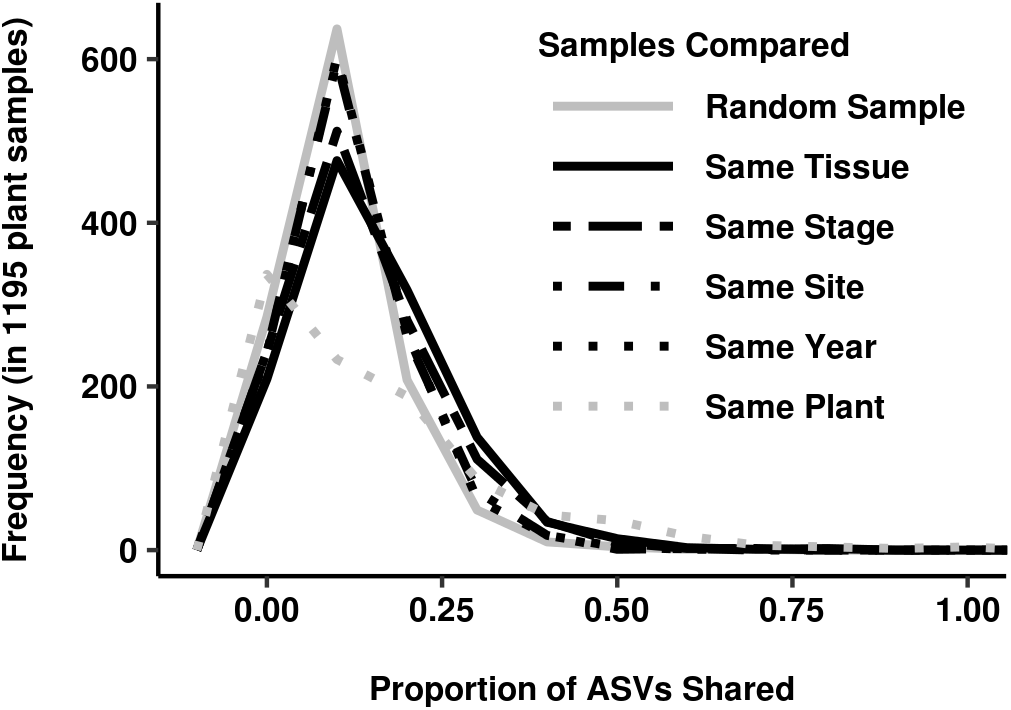
Bacterial assemblage composition was driven by the tissue sampled and the plant’s developmental stage at harvest. The frequency distribution for the proportion of ASVs shared by plant samples selected randomly (solid gray, median 0.096) differed from those for plant samples selected with respect to tissue type, developmental stage, site, or year (patterned black lines). The proportion of shared members was significantly higher when sample selection was conditioned on each variable, but the strongest shifts were observed for tissue type and developmental stage (same tissue median: 0.133, p < 2 × 10^-16^; same stage median: 0.117, p = 4 × 10^-11^; same site median: 0.106, p = 8 × 10^-5^; same year median: 0.100, p = 9 × 10^-4^). The frequency distribution for the proportion of ASVs shared with any samples taken from different tissues of the same individual plant is also shown (dotted gray line, p=0.038).

The influences of host and environment on assemblage composition were further supported by analysis of variance. After randomly subsampling to 1000 counts, composition variation between samples was quantified with respect to ASV presence by Raup-Crick dissimilarity (39), with respect to ASV abundance by Bray-Curtis dissimilarity (42), and with respect to ASV presence and phylogenetic relatedness by the unweighted UniFrac distance (43). Analysis of variance with permutation was performed on each dissimilarity matrix for each study variable (α = 0.001) (Supplementary Table S3). Variance between ecotypes was not greater than the variance within them, which is unsurprising given the genetic similarity of the haplogroup to which they belonged (40). Variance among plant individuals was significant only when compared by Bray-Curtis dissimilarity. Sample preparation plates and sequencing runs differed significantly in composition only when compared by Bray-Curtis dissimilarity and UniFrac distance. Tissues, developmental stages, planting sites, and years differed significantly in composition regardless of the dissimilarity or distance used to quantify differences.

Assemblages were primarily influenced by the type of host tissue sampled and its age. Study variables were nested according to the experimental design to create a multivariate model for permutational analysis of variance (PERMANOVA) (41). When PERMANOVA was performed with this model, tissue type and host developmental stage consistently explained the most sample variation regardless of the dissimilarity metric employed (Table 2). Tissue type was assigned between 13% and 38% of the total variance and developmental stage was assigned between 6% and 30% of variance. Since the residual sum-of-squares was markedly lowest in PERMANOVA on the Raup-Crick matrix, we focused on presence-absence variation in community composition when identifying the ASVs associated with specific tissues and developmental stages. An additional advantage of focusing on presence-absence variation is that 16S copy number varies among bacterial lineages.

**TABLE 2.** PERMANQVA results for rarefied data dissimilarity matrix ~ Year / Stage / Tissue + MiSeq Run / Sample Plate + Site

|  | DOF | Raup-Crick index |  |  | Bray-Curtis Dissimilarity |  |  | UniFrac Distance |  |  |
| --- | --- | --- | --- | --- | --- | --- | --- | --- | --- | --- |
|  |  | F | R<sup>2</sup> | Pr (>F) | F | R<sup>2</sup> | Pr (>F) | F | R<sup>2</sup> | Pr (>F) |
| Year : Stage : **Tissue** | 17 | 256.795 | 0.376 | < 0.001 | 7.330 | 0.127 | < 0.001 | 7.949 | 0.144 | < 0.001 |
| Year : **Stage** | 7 | 492.776 | 0.297 | < 0.001 | 12.233 | 0.087 | < 0.001 | 8.048 | 0.060 | < 0.001 |
| **Site** | 1 | 2275.626 | 0.196 | < 0.001 | 46.162 | 0.047 | < 0.001 | 26.216 | 0.028 | < 0.001 |
| **Year** | 1 | 1348.268 | 0.116 | < 0.001 | 28.543 | 0.029 | < 0.001 | 15.496 | 0.016 | < 0.001 |
| MiSeq Run : **Sample Plate** | 20 | 0 | 0 | 1 | 1.346 | 0.027 | < 0.001 | 1.389 | 0.030 | < 0.001 |
| **MiSeq Run** | 3 | 0 | 0 | 1 | 3.18 | 0.01 | < 0.001 | 5.59 | 0.018 | < 0.001 |
| Residuals |  |  | 0.015 |  |  | 0.673 |  |  | 0.704 |  |

### Assemblages in phyllosphere tissues became more distinguishable from those in roots as plants matured

Assemblages varied more between root and shoot tissues than within the phyllosphere. In principal coordinate analysis (PCoA) based on their dissimilarities (Figure 2A-C), samples from the stem and siliques clustered separately from rosette leaf samples and from root samples. This finding was robust to differences in rarefaction depth, filtering, and normalization of the count data (44) (Supplementary Figure S1). However, segregation along the first two principal coordinates was not clear when phyllosphere samples from flowering plants were ordinated alone, suggesting that most of the association with tissue type was driven by differences between root and shoot (Figure 2D-F). Supporting this interpretation, PERMANOVA on Raup-Crick dissimilarities of phyllosphere samples at flowering yielded a p-value below the significance threshold (α = 0.001) (Supplementary Table S4).

**Figure 2.**
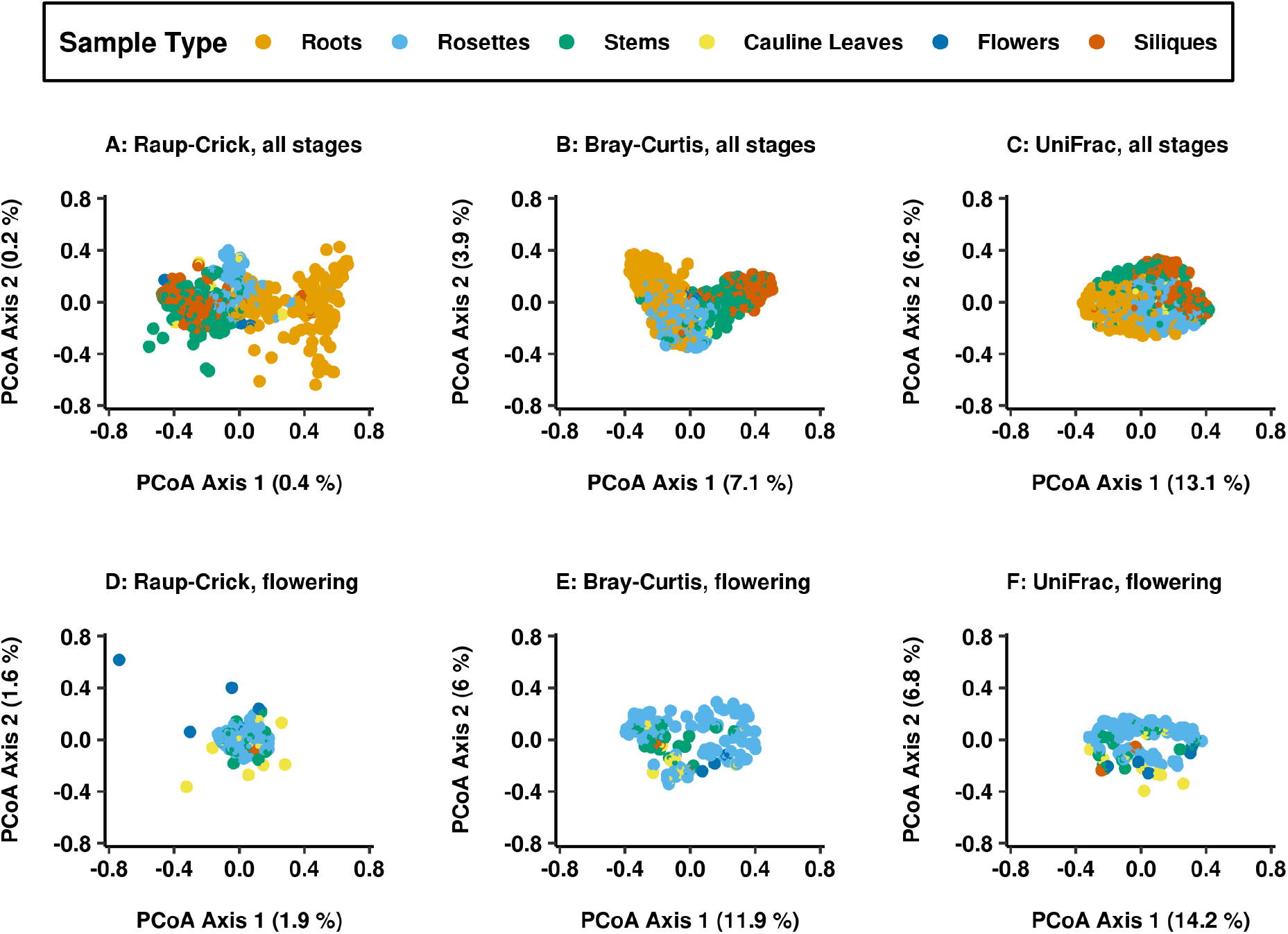
Roots and phyllosphere tissues housed distinct assemblages while tissues within the phyllosphere segregated between but not within stages. Plant samples segregated by tissue type in principal coordinate analysis (PCoA) based on their dissimilarities. The percentage of sample variance captured by the first two principal coordinates are listed on the x and y axis. (A,D) Raup-Crick dissimilarities are based on presence-absence differences between samples. (B,E) Bray-Curtis dissimilarities are based on quantitative differences in ASV counts between samples. (C,F) UniFrac distances incorporate phylogenetic relatedness of the ASVs present in samples based on the 16S gene tree. (A-C) For all dissimilarities, samples from roots (orange), rosette leaves (blue), and stems and siliques (green and red) clustered along the first coordinate. (D-F) In phyllosphere samples at flowering, rosette leaves (blue) overlapped with other phyllosphere tissues.

To disentangle the roles of tissue and age in defining habitats within the plant, we compared root and shoot tissues with respect to the ASVs present both before and after developmental transitions. Roots and rosettes were compared between late vegetative and flowering stages while roots and stems were compared between flowering and senescence. Tissue assemblages grew more distinguishable later in development, with the proportion of variance explained by tissue relative to other host variables increasing at later stages (Table 3). The differentiation of tissue habitats over time was further examined by quantifying their β diversity at each stage. Pairwise dissimilarities of samples within and between tissue types were calculated and the distributions of these distances were compared for both rosette leaves and roots (Figure 3). As development progressed, leaf assemblages simultaneously became more similar to each other and more distinct from those in the roots, perhaps due to unique selective pressures or a more restricted pool of potential colonists in the phyllosphere.

**TABLE 3.** PERMANOVA results for tissue comparisons at different stages

|  | Samples | Tissue R<sup>2</sup> | Tissue Pr (>F) |
| --- | --- | --- | --- |
| Vegetative Roots and Rosettes | 69 | 0.187 | 0.013 |
| Flowering Roots and Rosettes | 163 | 0.621 | < 0.001 |
| Flowering Roots and Stems | 94 | 0.391 | < 0.001 |
| Senescent Roots and Stems | 216 | 0.881 | < 0.001 |

**Figure 3.**
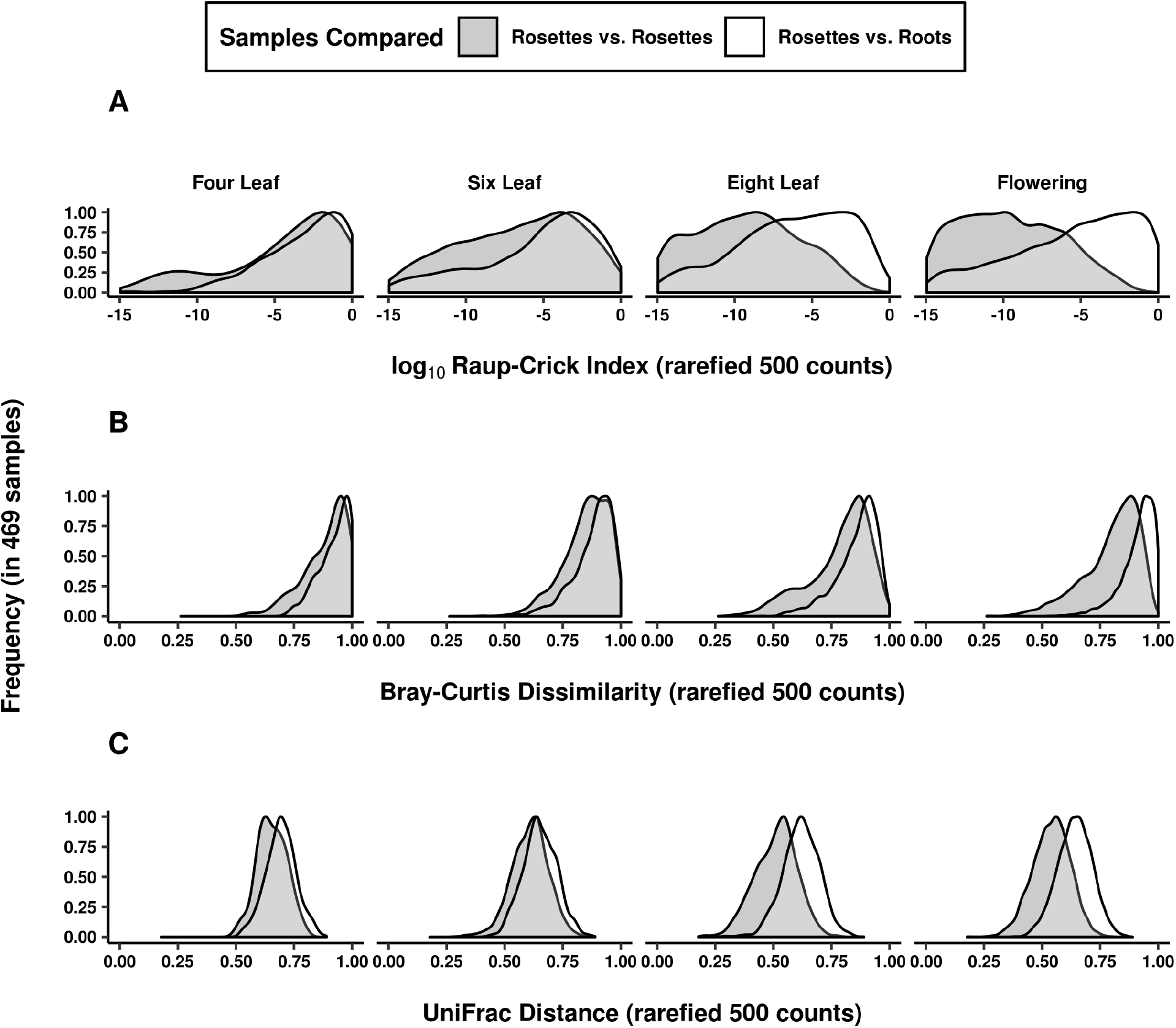
Phyllosphere assemblages became more distinguishable from those in roots as plants matured. The dataset was pruned to samples with at least 100 counts and 20 ASVs to calculate sample dissimilarities. The distribution of dissimilarities between rosette leaf samples (shaded distribution) was compared to the distribution of dissimilarities between rosette leaf and root samples (unshaded distributions). This procedure was repeated for (A) Raup-Crick dissimilarities, (B) Bray-Curtis dissimilarities, and (C) UniFrac distances. As plants matured (horizontal panels left to right), phyllosphere samples were increasingly similar to each other and distinguishable from the root samples.

### Assemblage associations with tissue and stage depended on sub-genus variation in bacteria

In the study of microbiome assembly, patterns can depend upon the taxonomic resolution with which bacteria are surveyed. ASVs represent the finest resolution of bacterial lineages possible with 16S data. If the effect of host tissue type was driven by differential filtering of closely related variants rather than larger bacterial clades, then it should weaken as taxonomic resolution decreases. To determine the dependence of the observed associations on taxonomic grouping, PERMANOVA was repeated with a Raup-Crick dissimilarity matrix produced with tables of ASV counts aggregated at the level of genus, family, order, class, and phylum (Table 4).

**TABLE 4.** PERMANOVA results for different taxonomic groupings

| Taxonomic Rank | % ASVs unassigned | Tissue |  | Stage |  |
| --- | --- | --- | --- | --- | --- |
|  |  | R<sup>2</sup> | Pr (>F) | R<sup>2</sup> | Pr (>F) |
| ASV | 0 | 0.803 | < 0.001 | 0.461 | < 0.001 |
| Genus | 43.94 | 0.769 | < 0.001 | 0.209 | 0.002 |
| Family | 17.71 | 0.682 | < 0.001 | 0.146 | 0.132 |
| Order | 8.17 | 0.202 | 0.003 | 0.155 | 0.004 |
| Class | 2.97 | 0.172 | < 0.001 | 0.081 | 0.065 |
| Phylum | 0.57 | 0.248 | < 0.001 | 0.153 | 0.001 |

Associations between assemblage composition and host tissue and age weakened with coarser taxonomic groupings of ASVs from genera to phylum. The weakening associations with coarser grouping suggested that the distribution of colonists within the host was not driven by widely shared bacterial traits but rather by variation within genera, whether acquired by horizontal transfer or evolved in vertically inherited traits. As a consequence, differences in the colonization patterns across tissues and stages are erased upon averaging the prevalence patterns of recently diverged lineages across taxonomic groups.

### Assemblage associations with tissue and stage were largely driven by the same ASVs, which were neither tissue-specific nor transient

By sampling different tissues at multiple timepoints, we were able to compare the bacterial lineages that distinguished assemblages across space and time. Spatial and temporal trends in the data were largely driven by the same colonists. To find ASVs significantly associated with specific tissues and developmental stages (α = 0.01), we compared the indicator value index of each ASV to a distribution generated by randomly permuting its presence-absence table (45). We found 460 ASVs that were significantly associated with specific tissue types; 70% of these were associated with roots and the remainder were associated with different phyllosphere tissues. Of the 268 stage-discriminating ASVs, 76% were also among the 460 that distinguished tissue types.

Assemblage differences between tissues were not driven by specialists and assemblage differences over time were not driven by transient community members. The ASVs associated with a specific tissue generally appeared in multiple plant tissues taken from a site in a given year rather than being restricted to a single habitat within the plant (93%). The ASVs associated with a specific stage generally recurred in a tissue throughout development rather than appearing at a single harvest stage (91%). Thus, assemblages differences did not result from the exclusive presence of ASVs in specific tissues or at specific stages in development, but rather from quantitative differences in prevalence over space and time.

### About a fifth of ASVs had consistent prevalence patterns across field sites and years while the rest had inconsistent spatial and temporal distributions

If tissue-specific host traits created environments favorable to particular colonists, then the spatial distributions of those colonists within the plant should be repeated across the sites and years in which they were observed. To assess whether ASV spatial distributions were repeated, the tissues in which ASVs reached maximum prevalence, when present, were compared between sites and years. Based on these comparisons, 21% (98/460) of the ASVs distinguishing tissues displayed consistent spatial trends, always peaking in prevalence in the same tissues (shown for Proteobacteria in Figure 4). For this fraction of ASVs, colonization patterns might be linked to tissue-specific host traits that differentially filter bacterial colonists. Of these consistently distributed ASVs, 79% were always most prevalent within roots, 11% in rosettes, 5% in stems, and 5% in siliques. Notably, the genus *Massilia* includes two distinct sets of ASVs that consistently peaked in different tissues; one set consistently peaked in roots while the second consistently peaked in siliques, emphasizing that sub-genus variation between bacterial lineages influenced their distributions within plants. For the remaining ASVs, including those in notable pathogen genera (Figure 5), spatial prevalence patterns were inconsistent between sites and years.

**Figure 4.**
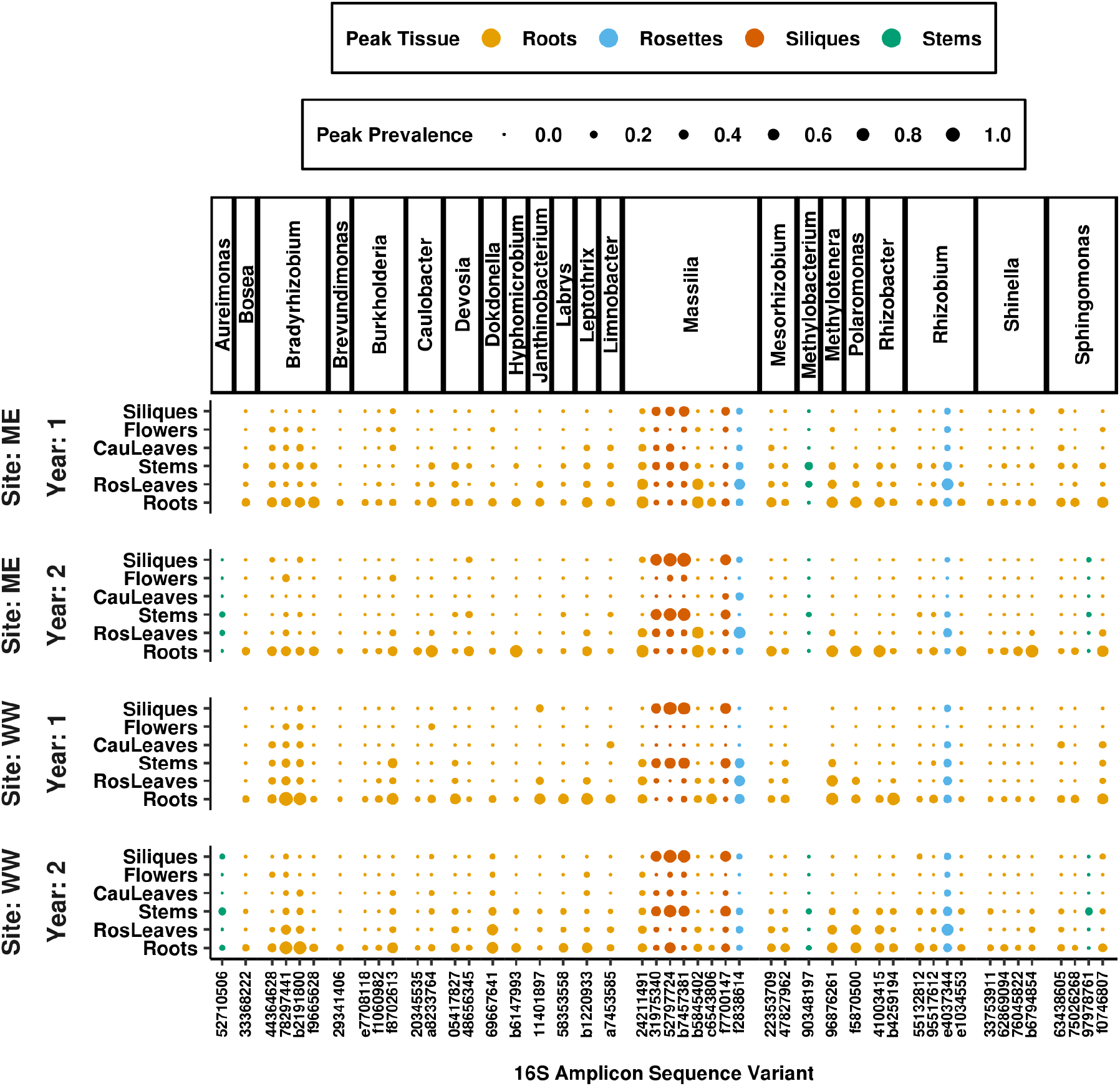
A fraction of ASVs behaved consistently across tissues at each site and in each year of the study. The ASVs distinguishing tissues, selected by indicator value indices, were filtered for those that reached maximum prevalence in the same tissue at each site and in each year when they were present. Proteobacteria ASVs with a total prevalence above five percent in all site and year combinations were selected for the plot. Panels on the y-axis separate sites and years and ASVs are grouped on the x-axis by genus. Dot sizes represent the maximum prevalence for each ASV in each tissue. Despite the significant association detected between assemblage composition and tissue type, only 21% of tissue-discriminating ASVs consistently reached peak prevalence in the same tissue. Of the ASVs that behaved consistently, 79% always reached peak prevalence in roots.

**Figure 5.**
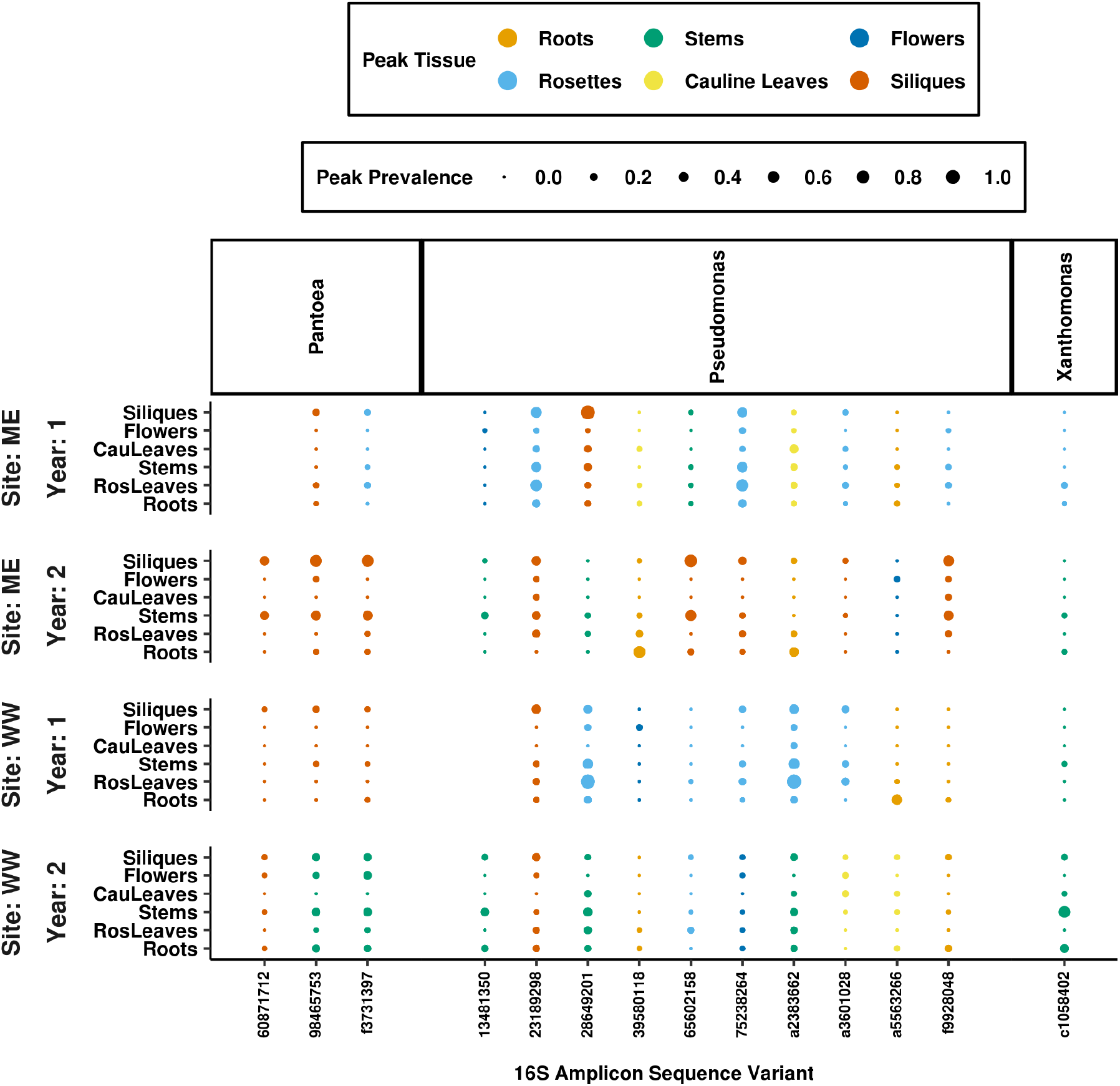
Most ASVs, including those in genera with pathogenic potential, do not have consistent spatial distributions across sites and years. The majority of ASVs distinguishing tissues did not reach maximum prevalence in the same tissue at each site and in each year when they were present. This pattern is exemplified by ASVs in the genera Pseudomonas, Xanthomonas, and Pantoea, which are notable for their pathogenic potential. Panels on the y-axis separate sites and years and ASVs are grouped on the x-axis by genus. Dot sizes represent the maximum prevalence for each ASV in each tissue.

Temporal distributions, like spatial ones, were highly variable. Of the ASVs driving associations with stage, only 23% (62/268) reached peak prevalence at a consistent plant developmental stage across sites and years. Temporal distributions were also dependent on fine taxonomic variation because ASVs within each genera displayed a variety of dynamics (Figure 6A). Among the Proteobacteria ASVs driving associations with age, the ones demonstrating the biggest changes in prevalence were found in both roots and rosettes across sites and years (Figure 6B). Temporally dynamic *Massilia* ASVs peaked during vegetative growth or flowering and then declined. Temporally dynamic *Methylobacterium* ASVs consistently increased during plant growth and peaked at senescence.

**Figure 6.**
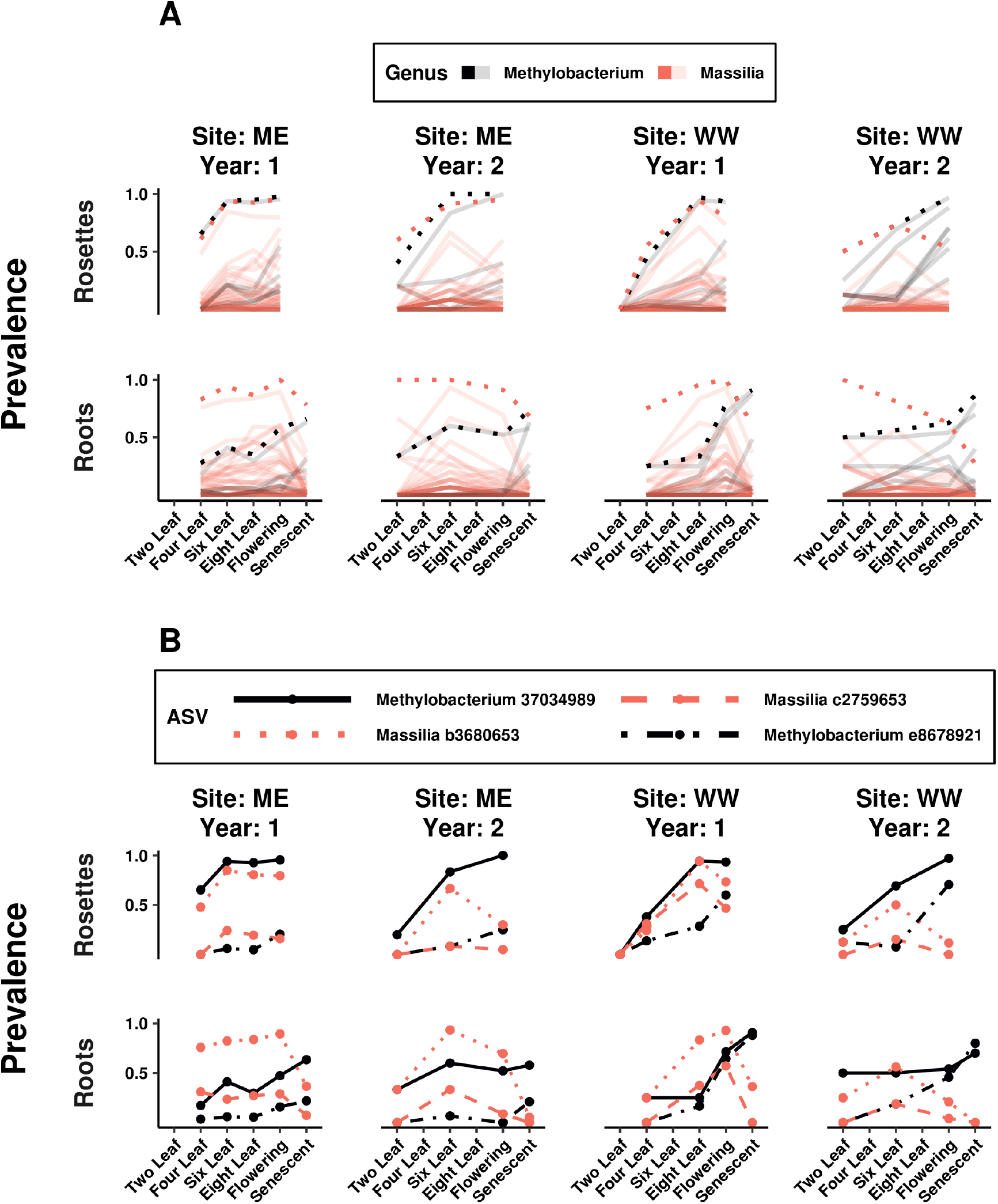
ASVs in the same genus have distinct temporal trends; a small number consistently reach high prevalence across sites and years despite variation in temporal trends. Plots feature Proteobacteria genera (*Massilia* in red and *Methylobacterium* in black) containing ASVs that distinguished developmental stages, based on indicator value indices, and reached over 70% prevalence in multiple sites and years. Horizontal panels separate the trends for roots and rosettes and vertical panels separate the trends for each site and year. (A) Bold trend lines show the temporal trends for counts grouped by genus, while transparent lines show the trends for individual ASVs in each genus. Genus-level trend lines subsume distinct ASV patterns. (B) Temporal colonization patterns were highly inconsistent despite the significant association between composition and stage, with only 23% of stagediscriminating ASVs consistently peaking at the same stage. Despite this variation, specific *Massilia* and *Methylobacterium* ASVs were among the most prevalent at both sites and in both years of study.

### Assemblages became more phylogenetically diverse and even over time

Temporal changes in assemblage α diversity can help explain the increased differentiation between assemblages inhabiting different tissues. The phylogenetic diversity of colonists was higher later in plant development (Figure 7A). Phylogenetic diversity was quantified by taking the tree of 16S variants present in each assemblage, weighting the branch lengths by variant abundance, and summing the branch lengths. This diversity trend was observed in each type of plant tissue sampled (Supplementary Figures S2) but not in samples of the surrounding soil, indicating it was related to the colonization of plant tissue and not purely driven by the abiotic environment during sample collection. The increasing phylogenetic diversity suggested that bacteria from across the tree had dispersed more widely among plant assemblages later in development. With more opportunities to encounter plants over time, subtle differences in the ASV colonization success between tissues were more likely to be exposed.

**Figure 7.**
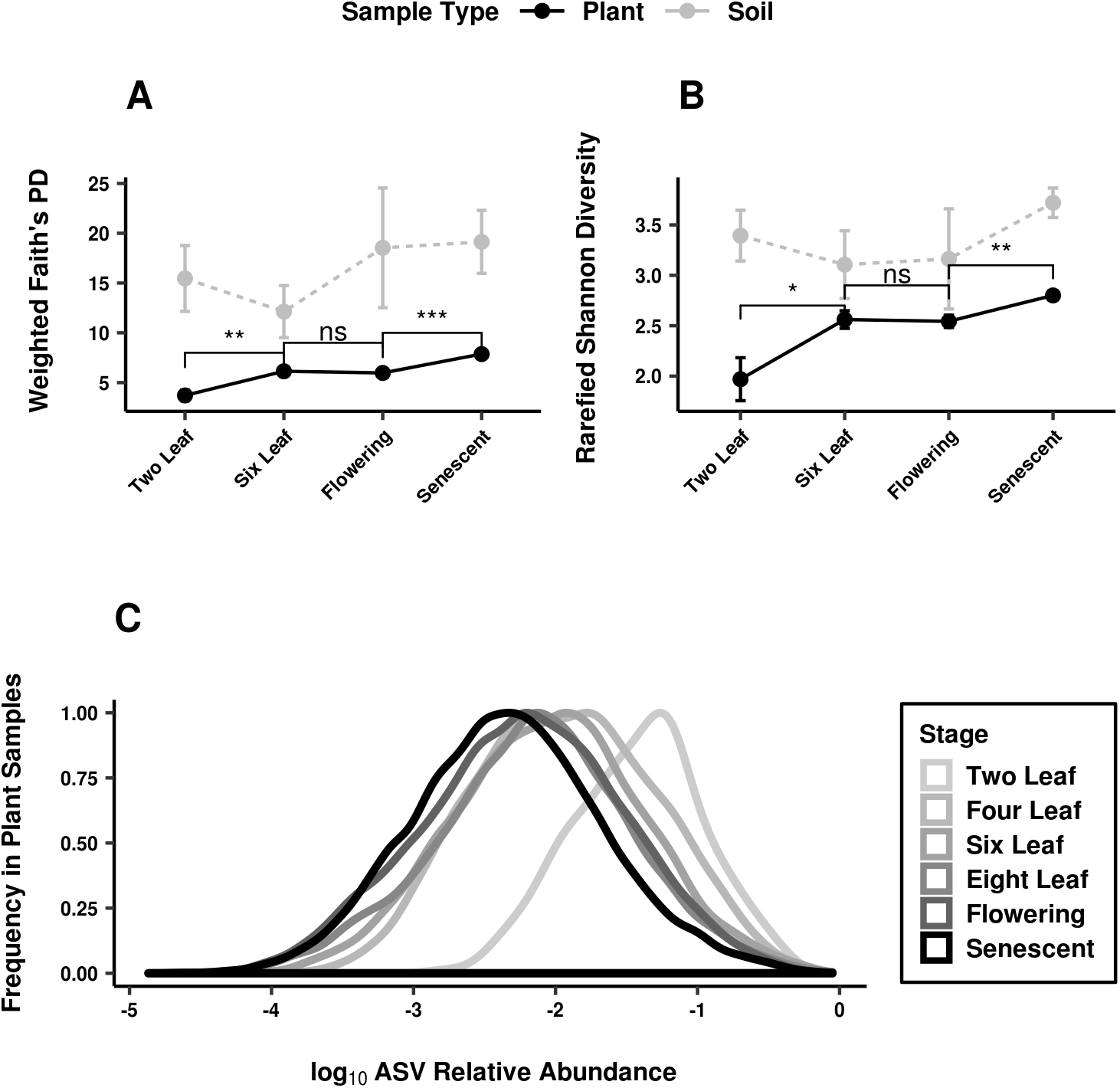
Assemblages within plants became more phylogenetically diverse and even over time. (A-B) Plots show mean and standard error for diversity measures of the plant (black) and soil (gray) samples at each developmental stage in the second year of study. Significance is shown for pairwise Wilcoxon tests between stages as follows: ns (not significant), * (p < 0.05), ** (p < 0.01), *** (p < 0.001), **** (p < 0.0001). (A) Phylogenetic distance was measured as the branch length on the 16S phylogenetic tree between ASVs in the sample, weighted by ASV abundances. It increased on average with developmental stage. (B) Shannon-Wiener (H’) indices from rarefied samples depended upon both the richness and evenness of samples and increased on average with developmental stage. (C) ASV relative abundances decrease during development as assemblages become more even. For plant samples with at least 100 counts, the relative abundance of each ASV present was calculated. The frequency distributions of these relative abundances are plotted for each developmental stage (shade), with relative abundance scaled by log10.

Temporal trends in assemblage evenness suggested that tissue type was related to the size of the colonization bottleneck for specific ASVs rather than to their ability to dominate other colonists once established in the endosphere. Variants that reached high prevalence in a tissue did not overtake assemblages in terms of relative abundance. The average Shannon-Wiener index (47) of assemblages increased (Figure 7B) and the distribution of ASV relative abundances decreased (Figure 7C) during plant life. Together, these trends showed that instead of domination by a small number of successful variants, mature tissues on average housed a more evenly represented set of ASVs.

## DISCUSSION

We surveyed bacterial assemblages inside roots and phyllosphere tissues throughout the life cycle of the model annual plant *A. thaliana*. Consistent with previous results, we found that assemblage composition differed between tissues (29) and during the course of development (20, 30). Because we examined multiple tissues and developmental stages in the same study, we were able to find three connections between these spatial and temporal trends in natural colonization that have not previously been identified.

First, the associations of host tissue type and developmental stage with assemblage composition were largely driven by the same colonists. These bacterial variants did not constitute omnipresent cores of tissue-specific inhabitants: those most strongly associated with tissue type typically did not reach even fifty percent prevalence within samples from a tissue type at a given time, underscoring the variability of community composition. Nor were these variants exclusive to specific habitat patch within the plant: most were observed at multiple sampling times and in multiple tissues. Instead, differential filtering over space and time quantitatively affected the prevalence of common endophytes, in agreement with a recent report of consistent organ occupancy in the plant microbiome (24).

Second, endophytic assemblages filtered by root and shoot tissues became more distinguishable later in development. Specifically, leaf assemblages grew more differentiated from root assemblages on average. ASVs with consistently higher prevalence in specific tissues suggested the existence of subtle differences between variants in the probability of colonizing different tissue niches. Increasing phylogenetic diversity indicated that bacterial lineages became more widely dispersed during plant development. As variants had more opportunities to colonize plant tissue, differences in the probability of colonizing particular tissues became more important in determining assemblage composition than the stochasticity of early colonization. Supporting this idea of the host plant gaining influence during assembly, a recent study of rice (*Oryza sativa*) roots found that microbiome composition was dynamic during vegetative growth and stabilized later in plant life (48).

A third major pattern in our data is the variability of spatiotemporal distributions among lineages within the same bacterial family or even genus. Most ASVs associated with tissue type (70%) were significantly enriched or depleted in the roots. Indeed, many of these ASVs belonged to genera that were consistently detected as root endophytes in a study of 17 European sites over 3 years (49). These included members of *Massilia, Burkholderia*, and *Bradyrhizobium*. However, *Massilia* also contained ASVs that reached peak prevalence in rosette leaves or siliques. An important factor in detecting associations between host and colonists is the resolution with which bacterial lineages are grouped in count tables. When microbes derived from natural sources, including plant leaves, are passaged outside the host in minimal media, they produce communities that are similar at the family level despite being highly variable at the level of sequence variants (50). In our study, grouping variants by family weakened associations with tissue type and erased the observed associations with harvest stage. Unlike the carbon metabolism traits that determined community structure *ex situ* (50), the functions selected by different tissue habitats may therefore not be shared broadly by lineages in a family.

In contrast to the roots, the phyllosphere presents microbes with a variety of challenges related to desiccation, toxins from other colonists, and motility (51). Adaptations known to mitigate these challenges, such as chemosensory and antimicrobial resistance genes, have occurred recently and vary at the sub-genus level in the lineages of known phyllosphere colonists (52) (53). Traits like these could lead to greater success in colonizing the leaf niche, creating phyllosphere assemblages that are more similar to each other and distant from root assemblages. As a result, closely related strains of known pathogenic, plant-beneficial, or biocontrol taxa might not establish in the same way throughout their plant hosts. Functional profiling of the bacterial assemblages may therefore be more valuable than taxonomic profiling in understanding their spatial and temporal trends within plants.

Our experimental design, including replicate sites and study years, allowed us to characterize consistency in distributions over space and time for the variants associated with host plant features. Biotic factors that can differ between roots and the phyllosphere, like salicylic acid, have been manipulated and shown to influence community assembly in both lab and field conditions (54,55). If such deterministic factors drove the observed variation in composition, then we might expect to see bacterial endophytes with the same across-tissue spatial distributions in repeated surveys. However, only about 20% of variants distinguishing tissues were consistently more prevalent in a specific tissue. The inconsistent behavior of most variants is perhaps not surprising given that samples typically did not share more than 15% of their colonists. This variability in the detection of endosphere colonists suggests that in nature, as in more controlled environments (17), stochastic factors drive much of community assembly. Even if individual lineages interact consistently with host plants, chance entry of functionally redundant strains from a large pool of soil colonists may give rise to mosaic assemblages.

Despite the growing strength of tissue effects during development, assemblages did not collapse to a few successful inhabitants of each tissue. In contrast, assemblage evenness increased in all plant tissues, agreeing with previous reports that community diversity increases as host plant tissues age (30,31). While host tissues appear to create different colonization bottlenecks for bacterial lineages, they did not appear to favor the dominance of these ASVs over others within assemblages.

These findings have consequences for the study of another key determinant of plant microbial communities: host genotype. Host genotype effects on colonization efficiency and microbiome composition are found both among angiosperm species that diverged hundreds of millions of years ago and among crop accessions that diverged through domestication within ten thousand years (19, 26, 56, 57, 58, 59). Compositional differences in the field can even be related to polymorphisms within a plant species (4,60). Since the variants associated with host variables are typically present throughout the plant and recurrent during development, filtering complex natural assemblages with these criteria can increase signal in the search for host polymorphisms linked to colonization success. Since tissue-associated assemblages become more distinct later in development, the host polymorphisms linked to variant prevalence or abundance might depend on which part of the plant is sampled, and when.

The effects of plant tissue type and developmental stage on assemblage composition were large compared to that of geographic site. Our results add to a growing body of evidence that the tissue sampled from a plant can explain more variation in microbial communities than geography. Studies of *A. thaliana* and *Boechera stricta* find that geographic sites and soil inocula play a substantial role in filtering plant microbial communities (20, 23). However, studies of cultivated *Agave* find that tissue explains more variation in community composition than species and site (27). Tissue type also explains more variation in epiphytic community composition than sample site in the species *S. taccada* (28). Together, these results suggest that plants are best considered as collections of distinct habitat patches for bacterial colonization.

## MATERIALS AND METHODS

### Planting

Field experiments were replicated over two years (2012-14) at two locations: Michigan State Southwest Michigan Research and Extension Center (ME) and University of Chicago Warren Woods Ecological Field Station (WW). Prior to planting in October, fields were tilled and grids were created with bottomless plastic pots (6-12 cm across) placed 2-5 cm into the ground and 10-30 cm apart. Within each grid, seeds for seven midwestern *A. thaliana* ecotypes were sown randomly and a fraction of pots were left empty for soil sampling. Seeds were surface-sterilized with ethanol and seedlings were thinned after germination with sterilized tweezers.

### Sample collection

Plant sampling order was randomized and all tools were flame-sterilized with ethanol between samples. Root and above-ground tissues were separated into sterile plastic tubes. For soil samples, sterile tubes were pushed 2-5cm into the ground. Tubes were stored at −80 C until processing. To remove loosely associated microbes, each plant sample was washed twice by vortexing with surfactant buffer (22). Plant samples were then transferred to Matrix tubes (Thermo Scientific, Waltham, MA, USA). Above-ground tissue was first separated into compartments with a scalpel and tweezers. For large tissues, only enough material was added to allow for bead homogenization. For soil, samples were put through a 2 mm sieve and ~100 mg was transferred to a Matrix tube. The tubes were randomized in 96-well racks with respect to sampling site, year, and timepoint. To dry the material, tubes were frozen to −80 C and lyophilized overnight. To powder the tissue, sterile silica beads were sealed into each tube with a SepraSeal cap (Thermo Scientific) and tubes were shaken on the 2010 Genogrinder (SPEX, Metuchen, NJ, USA) (1750 RPM, 2 min). Dry mass was recorded and up to 36 mg of material was retained per tube. All tubes were then randomized in Nunc 96-deepwell plates (Thermo Scientific) for DNA extraction.

### DNA extraction

Ground material was resuspended in TES (10 mM Tris-Cl, 1 mM EDTA, 100 mM NaCl) to a concentration of 0.04 mg/μL. Material was homogenized with the Genogrinder (1750 RPM) and homogenates (240 μL) were incubated (30 min) in new plates with lysozyme solution (Epicentre, Madison, WI, USA) at a final concentration of 50 U/μL. Proteinase-K (EMD Millipore, Billerica, MA, USA) and SDS were added to final concentrations of 0.5 mg/mL and 1%, respectively. Plates were incubated at 55 C for 4 h. An equal volume of 24:1 chloroform:isoamyl alcohol was mixed by pipette in each well. Plates were centrifuged at 6600xg with the Beckman Coulter Avanti J-25 (Beckman Instruments, Munich, Germany) for 15 min at 4 C. The top aqueous layer (350 μL) was removed and added to new plates with 500 μL 100% isopropanol. Plates were inverted to mix and incubated 1h at −20 C. After centrifugation for 15 min at 4 C, isopropanol was removed and DNA pellets were washed with 500 μL 70% ethanol. Pellets were dried in a chemical hood and resuspended in TE (100 μL, 10 mM Tris-Cl, 1 mM EDTA) by shaking. After incubation on ice for 5 min, plates were centrifuged for 12 min at 4 C and supernatants diluted 10X in TE were added to new 0.5 mL plates for PCR amplification.

### 16S rRNA gene amplification

The V5, V6, and V7 regions of the 16S rRNA gene were amplified from each sample using the 799F and 1193R primers with Illumina MiSeq adapters, and custom pads, linkers, and barcode sequences (61). The PCR volume was 25 μL: 1 μL of 10X diluted DNA template, 0.2 μM of each primer, 1X 5PRIME HotMasterMix (5PRIME, Gaithersburg, MD, USA), and 0.8X SBT-PAR additive (62). PCR amplification consisted of initial denaturation at 94 C for 2min, followed by 35 cycles of denaturation at 94 C for 30 s, annealing at 54.3 C for 40 s, and elongation at 68 C for 40 s, followed by a final elongation at 68 C for 7min. Each PCR was completed in triplicate, pooled, and purified with an equal volume of Axygen AxyPrep Mag PCR Clean-Up bead solution (Corning, Tewksbury, MA, USA). Amplicon concentrations were quantified by fluorimetry (QUANT-iT PicoGreen dsDNA Assay Kit, Life Technologies, Carlsbad, CA, USA) and 30 ng or a maximum of 30 μL per sample were pooled for six plates per sequencing run. Primer dimers and mitochondrial amplicons were removed by concentrating each amplicon pool 20X (Savant SPD121P SpeedVac Concentrator, Thermo Scientific) and purifying 300-700bp product with BluePippin (Sage Science, Beverly, MA, USA).

### Sequence data

Amplicon pools were sequenced using the Illumina MiSeq platform and MiSeq V2 Reagent Kits (Illumina, San Diego, CA, USA) to produce paired-end 250 bp reads (MiSeq Control Software v2.5.0.5). MiSeq Reporter v2.5.1.3 demultiplexed samples and removed reads without an index or matching PhiX. Within QIIME2, cutadapt removed primers from the paired reads and DADA2 identified ASVs. Primers 799F and 1193R were used to extract reads in silico from the QIIME-SILVA 16S database. These reads were used to build a classifier using QIIME2’s naive-bayes method and the sklearn algorithm was used to generate taxonomy assignments for the sequence variants. These assignments were used to filter any remaining mitochondrial and chloroplast sequences. Sequence variants with a frequency lower than 2 counts, samples with fewer than 10 reads, and samples with notes on irregularities during collection were also removed. To generate a phylogeny for the sequence variants, QIIME2 was used to align the sequences with MAFFT and to infer and root a phylogenetic tree. The tree was imported along with the DADA2-generated ASV count table, the taxonomy, and the metadata into a phyloseq (63) class in R (version 3.4.4) (64) for analysis. Count table transformations, pruning, and rarefaction were performed with phyloseq and distance matrix calculation, ordination, and PERMANOVA tests were performed with the vegan package (65). Phylogenetic analysis was performed with ape and picante (66,67). Figures and supplemental figures were produced with ggplot2, and ggpubr (68,69).

### Statistics

Three dissimilarity metrics were used to capture different aspects of microbiome variation. Presence-absence variation was represented by the Raup-Crick dissimilarity index, a probability of samples differing in composition based on ASV frequencies in the dataset. Alternatively, the Bray-Curtis dissimilarity quantified the abundance differences between ASV counts in each sample. The UniFrac distance incorporated presence-absence variation as well as phylogenetic relatedness between the ASVs present in samples based on the 16S gene tree.

ASVs associated with specific tissues or developmental stages were identified using the signassoc function of the indicspecies package (45,70). This function calculated an indicator value index (IndVal) based on the product of two probabilities: (1) the probability that a sample belonged to a habitat given ASV presence and (2) the probability that an ASV was present if a sample was taken from a habitat. For the habitats defined by each variable (six tissues, six developmental stages, two sites, and two years), indices were calculated independently for each ASV. The null hypothesis that no relationship existed between ASVs and conditions was tested by comparing the empirical index with a distribution generated by randomly permuting the ASV presence-absence count table. A two-tail p-value was used to select ASVs that are significantly more or less frequently observed in sampled belonging to a given condition (α = 0.01).

## Data Availability

Raw sequencing data is available in the NCBI’s Sequence Read Archive, BioProject ID PRJNA607544. The ASV count table, 16S phylogenetic tree and taxonomy, and sample metadata are available with the R commands used for analysis here: https://github.com/krbeilsmith/KBMP2020_Microbes

## Acknowledgements

KB and MP were supported by the University of Chicago Biological Sciences Division and by NIH T32 GM07197. MP was supported by a Department of Education GAANN grant in ecology to Cathy Pfister. Research was supported by NIH grant R01 GM083068 to JB. Computing resources were provided by The Center for Research Informatics, funded by the Biological Sciences Division at the University of Chicago with additional funding provided by the Institute for Translational Medicine, CTSA grant number UL1 TR000430 from the National Institutes of Health.

We thank Dave Francis at the Michigan State Southwest Michigan Research and Extension Center for providing and prepping field plots. We thank Carlos Sahagun for assisting with planting and Timothy Morton, Benjamin Brachi, Talia Karasov, Manfred Ruddat, and Roderick Woolley for assistance with sampling. The DNA extraction and 16S amplicon sequencing protocols were developed in collaboration with Benjamin Brachi and Alison Anastasio provided experimental design advice. Members of the Bergelson lab and the Department of Ecology and Evolution at the University of Chicago, in particular Caroline Oldstone-Jackson and Brooke Weigel, provided valuable feedback during the data processing and analysis. We thank two anonymous reviewers for their corrections and suggestions for the manuscript.

## SUPPLEMENTARY MATERIAL + FIGURE CAPTIONS

**Table S1. *Arabidopsis thaliana* ecotypes planted in the study (from the HPG-1 haplogroup)** File: TableS1.pdf

**Table S2. Sample collections for the study**

File: TableS2.pdf

**Table S3. Variables tested for association with dissimilarity matrices by PERMANOVA**

File: TableS3.pdf

**Table S4. PERMANOVA results for flowering phyllosphere samples**

File: TableS4.pdf

**Text S1. Supplementary Methods**

File: TextS1.pdf

**Text S2. Plants hosted bacterial assemblages distinct from those in the surrounding soil.** File: TextS2.pdf

**Figure S1. Plant tissue was the variable most strongly associated with sample composition, regardless of how composition variation was quantified or how the count tables for samples were transformed or thresholded.** The relationships of host tissue type and stage to community composition were visualized by principal coordinate (PCoA) ordination of the samples based on dissimilarities or distances (Figure S1). Samples clustered primarily by the material sampled regardless of the dissimilarity or distance used. Observed tissue differences were robust to common methods for minimizing the effects of different count totals among samples. (A) Weighted UniFrac is influenced by the phylogenetic relationships of ASVs present in samples as well as quantitative differences in their abundances. This distance was calculated for a rarefied dataset. (B) Bray-Curtis dissimilarity is based on quantitative differences in ASV abundance across samples. This dissimilarity was calculated for relative abundances in all samples, without rarefaction. A variance-stabilizing transformation (C) or a log transformation (D) was performed on the count table before the Bray-Curtis dissimilarity was taken. Finally, the importance of host variables relative to site and year was robust to thresholding the ASV count table based on the prevalence or abundance. (E) ASVs were filtered for prevalence to include only those presence in more than three samples. (F) ASVs were filtered for abundance to include only those with a count total exceeding one thousand for all samples.

File: FigureS1.tiff

**Figure S2. Diversity and evenness increased during development in all plant tissues.** All plots show mean and standard error for diversity measures of the plant samples at each developmental stage in the second year of study. Flower and cauline leaves were only sampled at one developmental stage and thus do not have trend lines. (A) Phylogenetic distance was measured as the branch length on the 16S phylogenetic tree between ASVs in the sample, weighted by ASV abundances. It increased on average with developmental stage. (B) Shannon-Wiener (H’) indices from rarefied samples depended upon both the richness and evenness of samples and increased on average with developmental stage.

File: FigureS2.tiff

## Notes

https://github.com/krbeilsmith/KBMP2020_Microbes

https://www.ncbi.nlm.nih.gov/bioproject/PRJNA607544

### Competing Interest Statement

The authors have declared no competing interest.

### Summary of Updates

Inserted correct Figure 7.

## References (in order of appearance)

(1) Hardoim PR, van Overbeek LS, Berg G, Pirttilä AM, Compant S, Campisano A, Döring M, Sessitsch A. 2015. The Hidden World within Plants: Ecological and Evolutionary Considerations for Defining Functioning of Microbial Endophytes. Microbiol Mol Biol Rev. 79(3):293–320.

(2) Khare E, Mishra J, Arora NK. 2018. Multifaceted Interactions Between Endophytes and Plant: Developments and Prospects. Front Microbiol. 9:2732.

(3) Kowalchuk GA, Yergeau E, Leveau JH, Sessitsch A, Bailey M, Liu W. 2010. Plant-associated microbial communities. Environmental molecular microbiology, 131–148.

(4) Bergelson J, Mittelstrass J, Horton MW. 2019. Characterizing both bacteria and fungi improves understanding of the *Arabidopsis* root microbiome. Scientific reports 9, no. 1:24.

(5) Rodríguez H, Fraga R. 1999. Phosphate solubilizing bacteria and their role in plant growth promotion. Biotechnol Adv. 17(4–5): 319–339. doi:10.1016/S0734-9750(99)00014-2.

(6) Franche C, Lindström K, Elmerich C. 2009. Nitrogen-fixing bacteria associated with leguminous and non-leguminous plants. Plant and soil, 321(1-2), 35–59.

(7) Marulanda A, Azcón R, Chaumont F, Ruiz-Lozano JM, Aroca R. 2010. Regulation of plasma membrane aquaporins by inoculation with a *Bacillus megaterium* strain in maize (*Zea mays* L.) plants under unstressed and salt-stressed conditions. Planta, 232(2), 533–543.

(8) Fernandez O, Theocharis A, Bordiec S, Feil R, Jacquens L, Clément C, Fontaine F, Barka EA. 2012. *Burkholderia phytofirmans* PsJN acclimates grapevine to cold by modulating carbohydrate metabolism. Molecular Plant-Microbe Interactions 25(4): 496–504.

(9) Mercado-Blanco J, Rodriguez-Jurado D, Hervás A, and Jiménez-Diaz RM. 2004. Suppression of *Verticillium* wilt in olive planting stocks by root-associated fluorescent *Pseudomonas* spp. Biological Control 30(2): 474–486.

(10) Kloepper JW, Ryu C. 2006. Bacterial endophytes as elicitors of induced systemic resistance. In Microbial root endophytes, pp. 33–52. Springer, Berlin, Heidelberg, 2006.

(11) Berg G, Köberl M, Rybakova D, Müller H, Grosch R, Smalla K. 2017. Plant microbial diversity is suggested as the key to future biocontrol and health trends. FEMS microbiology ecology, 93(5).

(12) Laine AL, Hanski I. 2006. Large-scale spatial dynamics of a specialist plant pathogen in a fragmented landscape. Journal of Ecology, 94(1), pp.217–226.

(13) Kuiper I, Bloemberg GV, Lugtenberg B.J. 2001. Selection of a plant-bacterium pair as a novel tool for rhizostimulation of polycyclic aromatic hydrocarbon-degrading bacteria. Molecular Plant-Microbe Interactions, 14(10), pp.1197–1205.

(14) Silva HSA, Romeiro RDS, Mounteer, A. 2003. Development of a root colonization bioassay for rapid screening of rhizobacteria for potential biocontrol agents. Journal of Phytopathology, 151(1), pp.42–46.

(15) Obadia B, Güvener ZT, Zhang V, Ceja-Navarro JA, Brodie EL, William WJ, Ludington WB. 2017. Probabilistic invasion underlies natural gut microbiome stability. Current Biology, 27(13), pp.1999–2006.

(16) Lazzaro BP, Fox GM. 2017. Host–Microbe Interactions: Winning the Colonization Lottery. Current Biology, 27(13), pp.R642–R644.

(17) Maignien L, DeForce EA, Chafee ME, Eren AM, Simmons SL. 2014. Ecological succession and stochastic variation in the assembly of *Arabidopsis thaliana* phyllosphere communities. mBio, 5(1), pp.e00682–13.

(18) Fitzpatrick CR, Copeland J, Wang PW, Guttman DS, Kotanen PM, Johnson MT. 2018. Assembly and ecological function of the root microbiome across angiosperm plant species. Proceedings of the National Academy of Sciences, 115(6), E1157–E1165.

(19) Bulgarelli D, Garrido-Oter R, Münch PC, Weiman A, Dröge J, Pan Y, McHardy AC, Schulze-Lefert P. 2015. Structure and function of the bacterial root microbiota in wild and domesticated barley. Cell Host Microbe. 17(3):392–403.

(20) Bulgarelli D, Rott M, Schlaeppi K, van Themaat EVL, Ahmadinejad N, Assenza F, Rauf P, Huettel B, Reinhardt R, Schmelzer E, Peplies J. 2012. Revealing structure and assembly cues for *Arabidopsis* root-inhabiting bacterial microbiota. Nature, 488(7409), 91.

(21) Mighell K, Saltonstall K, Turner BL, Espinosa-Tasón J, Van Bael, SA. 2019. Abiotic and biotic drivers of endosymbiont community assembly in *Jatropha curcas*. Ecosphere, 10(11).

(22) Lundberg DS, Lebeis SL, Paredes SH, Yourstone S, Gehring J, Malfatti S, Tremblay J, Engelbrektson A, Kunin V, Del Rio TG, Edgar RC. 2012. Defining the core *Arabidopsis thaliana* root microbiome. Nature, 488(7409), p.86.

(23) Wagner MR, Lundberg DS, Tijana G, Tringe SG, Dangl JL, Mitchell-Olds T. 2016. Host genotype and age shape the leaf and root microbiomes of a wild perennial plant. Nature Communications, 7, 12151.

(24) Massoni J, Bortfeld-Miller M, Jardillier L, Salazar G, Sunagawa S, Vorholt, JA. 2019. Consistent host and organ occupancy of phyllosphere bacteria in a community of wild herbaceous plant species. The ISME Journal, pp.1–14.

(25) Grady KL, Sorensen JW, Stopnisek N, Guittar J, Shade A. 2019. Abundance-occupancy distributions to prioritize plant core microbiome membership. Current opinion in microbiology, 49, pp.50–58.

(26) Germaine K, Keogh E, Garcia-Cabellos G, Borremans B, Van Der Lelie D, Barac T, Oeyen L, Vangronsveld J, Moore FP, Moore ER, Campbell CD. 2004. Colonisation of poplar trees by gfp expressing bacterial endophytes. FEMS Microbiology Ecology, 48(1), pp.109–118.

(27) Coleman-Derr D, Desgarennes D, Fonseca-Garcia C, Gross S, Clingenpeel S, Woyke T, North G, Visel A, Partida-Martinez LP, Tringe SG. 2016. Plant compartment and biogeography affect microbiome composition in cultivated and native *Agave* species. New Phytologist, 209(2), pp.798–811.

(28) Amend AS, Cobian GM, Laruson AJ, Remple K, Tucker SJ, Poff KE, Antaky C, Boraks A, Jones CA, Kuehu D, Lensing BR. 2019. Phytobiomes are compositionally nested from the ground up. PeerJ, 7, p.e6609.

(29) Bodenhausen N, Horton MW, Bergelson J. 2013. Bacterial communities associated with the leaves and the roots of *Arabidopsis thaliana*. PloS one, 8(2), e56329.

(30) Chaparro JM, Badri DV, Vivanco JM. 2014. Rhizosphere microbiome assemblage is affected by plant development. The ISME journal, 8(4), 790.

(31) Leff JW, Del Tredici P, Friedman WE, Fierer N. 2015. Spatial structuring of bacterial communities within individual *Ginkgo biloba* trees. Environmental Microbiology, 17(7), 2352–2361.

(32) Beattie GA, Lindow SE. 1999. Bacterial colonization of leaves: a spectrum of strategies. Phytopathology, 89(5), pp.353–359.

(33) Brandl MT, Lindow SE. 1998. Contribution of indole-3-acetic acid production to the epiphytic fitness of *Erwinia herbicola*. Appl. Environ. Microbiol., 64(9), pp.3256–3263.

(34) Callahan BJ, McMurdie PJ, Rosen MJ, Han AW, Johnson AJA, Holmes SP. 2016. DADA2: high-resolution sample inference from Illumina amplicon data. Nature methods, 13(7), 581.

(35) Bolyen E, Rideout JR, Dillon MR, Bokulich NA, Abnet C, Al-Ghalith GA, Alexander H, Alm EJ, Arumugam M, Asnicar F, Bai Y, Bisanz JE, Bittinger K, Brejnrod A, Brislawn CJ, Brown CT, Callahan BJ, Caraballo-Rodríguez AM, Chase J, Cope E, Da Silva R, Dorrestein PC, Douglas GM, Durall DM, Duvallet C, Edwardson CF, Ernst M, Estaki M, Fouquier J, Gauglitz JM, Gibson DL, Gonzalez A, Gorlick K, Guo J, Hillmann B, Holmes S, Holste H, Huttenhower C, Huttley G, Janssen S, Jarmusch AK, Jiang L, Kaehler B, Kang KB, Keefe CR, Keim P, Kelley ST, Knights D, Koester I, Kosciolek T, Kreps J, Langille MG, Lee J, Ley R, Liu Y, Loftfield E, Lozupone C, Maher M, Marotz C, Martin BD, McDonald D, McIver LJ, Melnik AV, Metcalf JL, Morgan SC, Morton J, Naimey AT, Navas-Molina JA, Nothias LF, Orchanian SB, Pearson T, Peoples SL, Petras D, Preuss ML, Pruesse E, Rasmussen LB, Rivers A, Robeson, II MS, Rosenthal P, Segata N, Shaffer M, Shiffer A, Sinha R, Song SJ, Spear JR, Swafford AD, Thompson LR, Torres PJ, Trinh P, Tripathi A, Turnbaugh PJ, Ul-Hasan S, van der Hooft JJ, Vargas F, Vázquez-Baeza Y, Vogtmann E, von Hippel M, Walters W, Wan Y, Wang M, Warren J, Weber KC, Williamson CH, Willis AD, Xu ZZ, Zaneveld JR, Zhang Y, Zhu Q, Knight R, Caporaso JG. 2018. QIIME 2: Reproducible, interactive, scalable, and extensible microbiome data science. 6:e27295v2 https://doi.org/10.7287/peerj.preprints.27295v2

(36) Price MN, Dehal PS, Arkin AP. 2010. FastTree 2–approximately maximum-likelihood trees for large alignments. PloS one, 5(3), e9490.

(37) Katoh K, Standley DM. 2013. MAFFT multiple sequence alignment software version 7: improvements in performance and usability. Molecular biology and evolution, 30(4), 772–780.

(38) Quast C, Pruesse E, Yilmaz P, Gerken J, Schweer T, Yarza P, Peplies J, Glöckner FO. 2012. The SILVA ribosomal RNA gene database project: improved data processing and web-based tools. Nucleic acids research, 41(D1), D590–D596.

(39) Raup DM, Crick RE. 1979. Measurement of faunal similarity in paleontology. Journal of Paleontology, 1213–1227.

(40) Exposito-Alonso M, Becker C, Schuenemann VJ, Reiter E, Setzer C, Slovak R, Brachi B, Hagmann J, Grimm DG, Chen J, Busch W. 2018. The rate and potential relevance of new mutations in a colonizing plant lineage. PLoS genetics, 14(2), p.e1007155.

(41) Anderson MJ. 2014. Permutational multivariate analysis of variance (PERMANOVA). Wiley statsref: statistics reference online, 1–15.

(42) Bray JR, Curtis JT. 1957. An ordination of the upland forest communities of southern Wisconsin. Ecological monographs, 27(4), 325–349.

(43) Lozupone C, Knight R. 2005. UniFrac: a new phylogenetic method for comparing microbial communities. Appl. Environ. Microbiol., 71(12), 8228–8235.

(44) Weiss S, Xu ZZ, Peddada S, Amir A, Bittinger K, Gonzalez A, Lozupone C, Zaneveld JR, Vázquez-Baeza Y, Birmingham A, Hyde ER. 2017. Normalization and microbial differential abundance strategies depend upon data characteristics. Microbiome, 5(1), p.27.

(45) Cáceres MD, Legendre P. 2009. Associations between species and groups of sites: indices and statistical inference. Ecology, 90(12), pp.3566–3574.

(46) McCoy CO, Matsen IV FA. 2013. Abundance-weighted phylogenetic diversity measures distinguish microbial community states and are robust to sampling depth. PeerJ, 1, e157.

(47) Shannon CE. 1948. A mathematical theory of communication. Bell system technical journal, 27(3), 379–423.

(48) Edwards JA, Santos-Medellín CM, Liechty ZS, Nguyen B, Lurie E, Eason S, Phillips G, Sundaresan V. 2018. Compositional shifts in root-associated bacterial and archaeal microbiota track the plant life cycle in field-grown rice. PLoS biology, 16(2), p.e2003862.

(49) Thiergart T, Durán P, Ellis T, Vannier N, Garrido-Oter R, Kemen E, Roux F, Alonso-Blanco C, Ågren J, Schulze-Lefert P, Hacquard S. 2020. Root microbiota assembly and adaptive differentiation among European *Arabidopsis* populations. Nature Ecology & Evolution, 4(1), pp.122–131.

(50) Goldford JE, Lu N, Bajić D, Estrela S, Tikhonov M, Sanchez-Gorostiaga A, Segrè D, Mehta P, Sanchez A. 2018. Emergent simplicity in microbial community assembly. Science, 361(6401), pp.469–474.

(51) Vorholt JA. 2012. Microbial life in the phyllosphere. Nature Reviews Microbiology, 10(12), pp.828–840.

(52) Clarke CR, Hayes BW, Runde BJ, Markel E, Swingle BM, Vinatzer BA. 2016. Comparative genomics of *Pseudomonas syringae* pathovar tomato reveals novel chemotaxis pathways associated with motility and plant pathogenicity. PeerJ, 4, p.e2570.

(53) Hwang MS, Morgan RL, Sarkar SF, Wang PW, Guttman DS. 2005. Phylogenetic characterization of virulence and resistance phenotypes of *Pseudomonas syringae*. Appl. Environ. Microbiol., 71(9), pp.5182–5191.

(54) Lebeis SL, Paredes SH, Lundberg DS, Breakfield N, Gehring J, McDonald M, Malfatti S, Del Rio TG, Jones CD, Tringe SG, Dangl JL. 2015. Salicylic acid modulates colonization of the root microbiome by specific bacterial taxa. Science, 349 (6250), pp.860–864.

(55) Traw MB, Kniskern JM, Bergelson J. 2007. SAR increases fitness of *Arabidopsis thaliana* in the presence of natural bacterial pathogens. Evolution: International Journal of Organic Evolution, 61(10), pp.2444–2449.

(56) Kembel SW, O’Connor TK, Arnold HK, Hubbell SP, Wright SJ, Green JL. 2014. Relationships between phyllosphere bacterial communities and plant functional traits in a neotropical forest. Proceedings of the National Academy of Sciences, 111(38), 13715–13720.

(57) Laforest-Lapointe I, Messier C, Kembel SW. 2016. Tree phyllosphere bacterial communities: exploring the magnitude of intra-and inter-individual variation among host species. PeerJ, 4, e2367.

(58) Peiffer JA, Spor A, Koren O, Jin Z, Tringe SG, Dangl JL, Buckler ES, Ley RE. 2013. Diversity and heritability of the maize rhizosphere microbiome under field conditions. Proceedings of the National Academy of Sciences, 110(16), pp.6548–6553.

(59) Barak JD, Kramer LC, Hao LY. 2011. Colonization of tomato plants by *Salmonella enterica* is cultivar dependent, and type 1 trichomes are preferred colonization sites. Appl. Environ. Microbiol., 77(2), pp.498–504.

(60) Horton MW, Bodenhausen N, Beilsmith K, Meng D, Muegge BD, Subramanian S, Vetter MM, Vilhjálmsson BJ, Nordborg M, Gordon JI, Bergelson J. 2014. Genome-wide association study of *Arabidopsis thaliana* leaf microbial community. Nature Communications, 5, p.5320.

(61) Kozich JJ, Westcott SL, Baxter NT, Highlander SK, Schloss, PD. 2013. Development of a dual-index sequencing strategy and curation pipeline for analyzing amplicon sequence data on the MiSeq Illumina sequencing platform. Appl. Environ. Microbiol., 79(17), pp.5112–5120.

(62) Samarakoon T, Wang SY, Alford MH. 2013. Enhancing PCR amplification of DNA from recalcitrant plant specimens using a trehalose-based additive. Applications in Plant Sciences, 1(1), p.1200236.

(63) R Core Team. 2018. R: A language and environment for statistical computing. R Foundation for Statistical Computing, Vienna, Austria. https://www.R-project.org/.

(64) McMurdie PJ, Holmes S. 2013. phyloseq: an R package for reproducible interactive analysis and graphics of microbiome census data. PloS one, 8(4), p.e61217.

(65) Oksanen J, Blanchet FG, Friendly M, Kindt R, Legendre P, McGlinn D, Minchin PR, O’Hara RB, Simpson GL, Solymos P, Stevens MHH, Szoecs E, Wagner H. 2017. vegan: Community Ecology Package. R package version 2.4-5. https://CRAN.R-project.org/package=vegan

(66) Paradis E, Schliep K. 2018. ape 5.0: an environment for modern phylogenetics and evolutionary analyses in R. Bioinformatics, 35(3), pp.526–528.

(67) Kembel SW, Cowan PD, Helmus MR, Cornwell WK, Morlon H, Ackerly DD, Blomberg SP, Webb CO. 2010. Picante: R tools for integrating phylogenies and ecology. Bioinformatics 26:1463–1464.

(68) Wickham H. ggplot2: Elegant Graphics for Data Analysis. Springer-Verlag New York, 2016.

(69) Kassambara A (2018). ggpubr: ‘ggplot2’ Based Publication Ready Plots. R package version 0.1.8. https://CRAN.R-project.org/package=ggpubr

(70) Cáceres, M. D. (2020). How to use theindicspeciespackage (ver. 1.7.8) https://cran.r-project.org/web/packages/indicspecies/vignettes/indicspeciesTutorial.pdf

