## Supplementary material for "Natural bacterial assemblages in *Arabidopsis thaliana* tissues become more distinguishable and diverse during host development": Table S1

**TABLE S1** Arabidopsis thaliana ecotypes planted in the study (from the HPG-1 haplogroup)

| Accession | Ecotype ID | Collection Location | Coordinates |
|---|---|---|---|
| BRR4 | 470 | Watseka, IL, USA | (40.8313, - 87.735) |
| LI-WP-041 | 546 | Nissequogue, NY, USA | (40.9076, - 73.2089) |
| L-R-10 | 1797 | Union Pier, MI, USA | (41.847, - 86.67) |
| MNF-Che-47 | 1942 | Manistee National Forest, MI, USA | (43.5251, - 86.1843) |
| Pent-7 | 2191 | Pentwater, MI, USA | (43.7623, - 86.3929) |
| PT1.85 | 8057 | Hanna, IN, USA | (41.3432, - 86.7368) |
| SLSP-69 | 2285 | Silver Lake, MI, USA | (43.665, - 86.496) |
