## Supplementary material for "Natural bacterial assemblages in *Arabidopsis thaliana* tissues become more distinguishable and diverse during host development": Table S2

**TABLE S2** Sample collections for the study

| Year | Date | Site | Stage | Sample type(s) |
|---|---|---|---|---|
| 1 | 10/12/12 | ME | No Plant | Soil |
| 1 | 10/15/12 | WW | No Plant | Soil |
| 1 | 10/26/12 | ME | Two Leaf | Roots, Rosette Leaves |
| 1 | 10/29/12 | WW | Two Leaf | Roots, Rosette Leaves |
| 1 | 11/29/12 | ME,WW | Four Leaf | Roots, Rosette Leaves |
| 1 | 02/13/13 | ME | Six Leaf | Roots, Rosette Leaves |
| 1 | 03/29/13 | ME, WW | Eight Leaf | Roots, Rosette Leaves |
| 1 | 05/01/13 | ME, WW | Flowering | Roots, Rosette Leaves, Stems, Cauline Leaves, Flowers, Siliques |
| 1 | 06/14/13 | ME, WW | Senescent | Roots, Stems, Siliques |
| 2 | 10/28/13 | ME, WW | No Plant | Soil |
| 2 | 12/04/13 | ME, WW | Two Leaf | Soil, Roots, Rosette Leaves |
| 2 | 04/16/14 | ME, WW | Six Leaf | Soil, Roots, Rosette Leaves |
| 2 | 05/15/14 | ME, WW | Flowering | Soil, Roots, Rosette Leaves, Stems, Cauline Leaves, Flowers, Siliques |
| 2 | 07/03/14 | ME, WW | Senescent | Soil, Roots, Stems, Siliques |
