## Supplementary material for "Natural bacterial assemblages in *Arabidopsis thaliana* tissues become more distinguishable and diverse during host development": Table S3

**TABLES S3** Variables tested for association with dissimilarity index matrices by PERMANOVA

| | | | Raup-Crick index | | | | Bray-Curtis dissimilarity | | | UniFrac distance | | |
|---|---|---|---|---|---|---|---|---|---|---|---|---|
| Variable | Description | Values | DOF | F | $R^2$ | Pr (>F) | F | $R^2$ | Pr (>F) | F | $R^2$ | Pr (>F) |
| Tissue | The part of the plant washed, ground, and sampled for DNA extraction | Roots, Rosettes, Stems, Cauline Leaves, Flowers, Siliques | 5 | 577.184 | 0.803 | < 0.001 | 23.909 | 0.145 | < 0.001 | 21.600 | 0.133 | < 0.001 |
| Stage | The developmental stage of the plant when the tissue was harvested | Two Leaf, Four Leaf, Six Leaf, Eight Leaf, Flowering, Senescent | 5 | 120.622 | 0.461 | < 0.001 | 14.606 | 0.094 | < 0.001 | 9.091 | 0.060 | < 0.001 |
| Site | The location of the field where the plant grew | Michigan Extension, Warren Woods | 1 | 202.328 | 0.222 | < 0.001 | 38.890 | 0.052 | < 0.001 | 22.897 | 0.031 | < 0.001 |
| Year | The year in which the plant was harvested | 1, 2 | 1 | 93.356 | 0.116 | < 0.001 | 21.224 | 0.029 | < 0.001 | 11.904 | 0.016 | < 0.001 |
| Ecotype | The *A. thaliana* genotype of the plant sampled | BRR4, LI-WP-041, L-R-10, MNF-Che-47, Pent-7, PT1.85, SLSP-69 | 6 | 1.717 | 0.014 | 0.342 | 0.931 | 0.008 | 0.748 | 0.802 | 0.007 | 0.958 |
| Sample Plate | The 96-well microplate in which sample extraction and 16S amplification were performed | Plate 1 – 24 | 23 | 0 | 0 | 0.914 | 1.311 | 0.042 | < 0.001 | 1.640 | 0.052 | < 0.001 |
| MiSeq Run | The batch in which samples were sequenced on the Illumina MiSeq platform | Run 1 – 4 | 3 | 0 | 0 | 0.710 | 2.296 | 0.010 | < 0.001 | 4.200 | 0.017 | < 0.001 |
| Plant ID | The plant individual from which tissues were harvested | Plant 1 – 376 | 375 | 1.253 | 0.448 | 0.012 | 1.172 | 0.431 | < 0.001 | 1.067 | 0.408 | 0.003 |
