## Supplementary material for "Natural bacterial assemblages in *Arabidopsis thaliana* tissues become more distinguishable and diverse during host development": Table S4

**TABLE S4** PERMANOVA results for flowering phyllosphere samples

| | | Raup-Crick index | | |
|---|---|---|---|---|
| | DOF | F | $R^2$ | Pr (>F) |
| Year : **Tissue** | 7 | 11.385 | 0.380 | 0.01 |
| **Site** | 1 | 42.098 | 0.201 | < 0.001 |
| **Year** | 1 | 1.843 | 0.009 | 0.421 |
| MiSeq Run : **Sample Plate** | 20 | 0 | 0 | 0.917 |
| **MiSeq Run** | 3 | 0 | 0 | 0.773 |
| Residuals | | | 0.015 | |
