## Supplementary figures and images for "Natural bacterial assemblages in *Arabidopsis thaliana* tissues become more distinguishable and diverse during host development"

### Figure S1

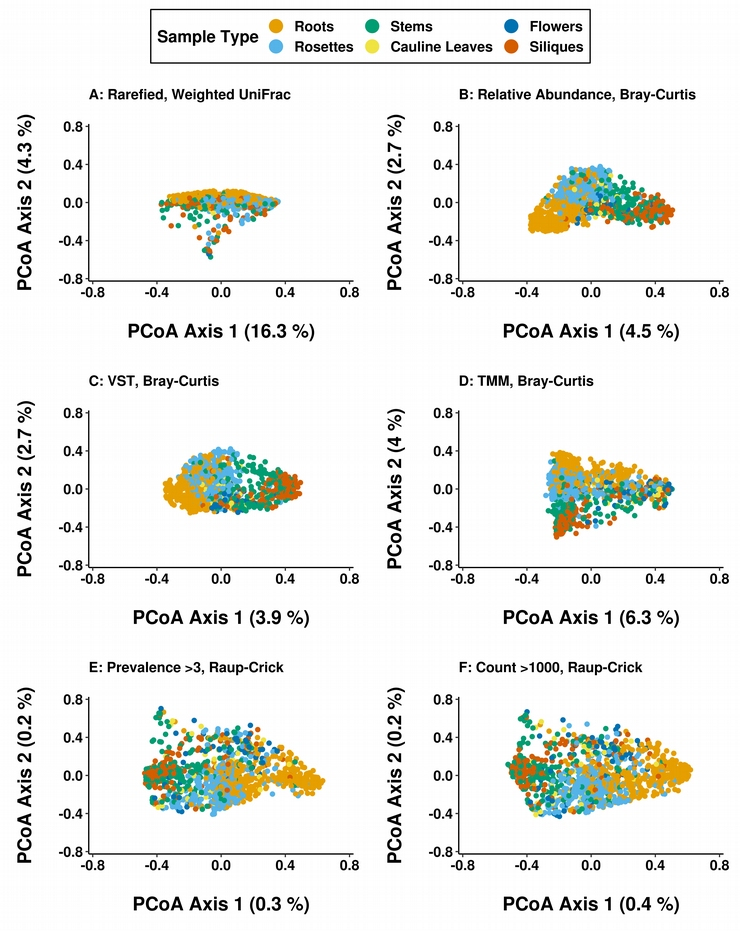

### Figure S2

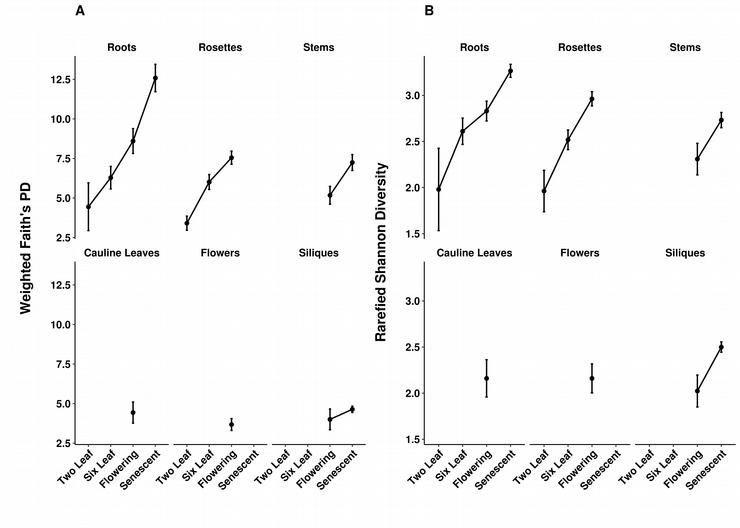
